# Engineered Red Blood Cells Carrying PCSK9 Inhibitors Persistently Lower LDL and Prevent Obesity

**DOI:** 10.1101/2020.11.02.364620

**Authors:** Rhogerry Deshycka, Valentino Sudaryo, Nai-Jia Huang, Yushu Xie, Liyan Y. Smeding, Moon Kyung Choi, Hidde L. Ploegh, Harvey F. Lodish, Novalia Pishesha

**Affiliations:** Department of Biological Engineering, Massachusetts Institute of Technology, Cambridge, MA; Whitehead Institute for Biomedical Research, Cambridge, MA; Program in Cellular and Molecular Medicine, Boston Children’s Hospital, Boston, MA; Brigham and Women’s Hospital, Boston, MA; Department of Biology, Massachusetts Institute of Technology, Cambridge, MA; Department of Pharmaceutical Chemistry, University of California, San Francisco, CA; Department of Biochemistry, Stanford University School of Medicine, Stanford, CA; Society of Fellows, Harvard University, Cambridge, MA

## Abstract

Low plasma levels of Proprotein Convertase Subtilisin/Kexin 9 (PCSK9) are associated with decreased low-density lipoprotein (LDL) cholesterol and a reduced risk of cardiovascular disease. PCSK9 binds to the epidermal growth factor-like repeat A (EGFA) domain of LDL receptors (LDLR), very low-density lipoprotein receptors (VLDLR), apolipoprotein E receptor 2 (ApoER2), and lipoprotein receptor–related protein 1 (LRP1) and accelerates their degradation, thus acting as a key regulator of lipid metabolism. Antibody and RNAi - based PCSK9 inhibitor treatments lower cholesterol and prevent cardiovascular incidents in patients, but their high cost hampers market penetration. We sought to develop a safe, long-term and one-time solution to treat hyperlipidemia. We created a cDNA encoding a chimeric protein in which the extracellular N-terminus of glycophorin A was fused to the LDLR EGFA domain and introduced this gene into mouse bone marrow hematopoietic stem and progenitor cells (HSPCs). Following transplantation into irradiated mice, the animals produced red blood cells (RBCs) with the EGFA domain (EGFA-GPA RBCs) displayed on their surface. These animals showed significantly reduced plasma PCSK9 (66.5% decrease) and reduced LDL levels (40% decrease) for as long as 12 months post-transplantation. Furthermore, the EGFA-GPA mice remained lean for life and maintained normal body weight under high-fat diet. Hematopoietic stem cell gene therapy can generate red blood cells expressing an EGFA - glycophorin A chimeric protein as a practical and long-term strategy for treating chronic hyperlipidemia and obesity.

## Introduction

Hyperlipidemia, characterized by elevated plasma LDL (low-density lipoprotein) levels, is a risk factor for cardiovascular disease, the leading cause of death worldwide. Therapeutics that reduce plasma LDL levels are the mainstay in the prevention of cardiovascular disease. Statins, small molecule inhibitors of HMG-CoA reductase, impede the generation of LDL cholesterol and are effective in reducing the incidence of cardiovascular disease and mortality in high risk individuals. Because statins inhibit a crucial metabolic pathway, patients who receive statin medication can suffer adverse effects, ranging from minor cognitive decline and muscle pain to an increased risk of diabetes as well as liver and kidney failure [1].

The discovery of proprotein convertase subtilisin-like kexin type 9 (PCSK9) was a possible breakthrough in tackling hypercholesterolemia. PCSK9 is produced by the liver and accelerates degradation of LDLR. A gain of function mutation in PCSK9 in two French families that exhibit a dominant form of Familial Hypercholesterolemia suggested that reducing PCSK9 levels is an attractive mechanism to lower cholesterol [2]. A woman of African descent who had a loss of function mutation in both PCSK9 alleles had significantly lower plasma LDL levels while otherwise healthy. Combined, these genetic findings suggested that inhibition of PCSK9 could be a novel and safe target to treat hypercholesterolemia [3].

PCSK9 is a serine protease encoded by a gene comprising 12 exons located on chromosome 1p32. PCSK9 binds to the epidermal growth factor-like A domain (EGFA) on the stalk of the LDLR and blocks a structural transition of LDLR in the endosome. Instead of recycling to the surface [4], LDLR is targeted for lysosomal degradation. Inhibitory peptides that mimic EGFA prevent degradation of LDLR and can lower plasma LDL levels [5]. The catalytic subunit of PCSK9 binds structurally similar EGFA domains on LDLR superfamily members including very low-density lipoprotein receptors (VLDLR), apolipoprotein E receptor 2 (ApoER2), and lipoprotein receptor-related protein 1 (LRP1), implicating PCSK9 as key regulator in lipid metabolism [6]. PCSK9 also regulates triglyceride metabolism by enhancing the degradation of CD36 (cluster of differentiation 36), a scavenger receptor involved in transport of long-chain fatty acids and triglyceride storage in adipocytes and liver [7]. Independent of its action on LDLR, PCSK9 modulates cholesterol transport and metabolism in the intestines by upregulating the intestinal epithelial cholesterol transporter NPC1L1 (Niemann-Pick C1-like protein 1) and increasing the expression of apolipoprotein B48 [8].

Multiple molecular strategies and therapeutics have been developed that inhibit LDLR degradation by PCSK9 and thus lower LDL levels (for review see reference [9]). While showing promise, these therapeutics have drawbacks, such as a transient reduction of LDL that must be counteracted by frequent injections of antibodies, peptides, or small molecule drugs. FDA-approved antibody-based drugs, while showing efficacy in patients, are expensive and limit their wide-use as a preventative strategy [10]. Disabling the PCSK9 gene using gene editing technologies may allow a one-shot injection as a preventative treatment to reduce the risk of cardiovascular disease [11]. Red blood cells are an attractive vehicle to introduce therapeutics into the body due to their systemic circulation, long half-life, and lack of DNA and RNA [12–14]. Hematopoietic stem cell (HSC) gene therapy – autologous transplantation of HSCs in which a cDNA is expressed by a lentivirus vector– has proven successful in treating X-Linked Severe Combined Immunodeficiency, Wiskott Aldrich Disease, Sickle Cell Anemia, and other conditions [15–18].

Here we generated a cDNA that encodes a chimeric protein in which the extracellular N-terminus of glycophorin A was fused to the LDLR EGFA domain. We introduced this gene into mouse bone marrow hematopoietic stem and progenitor cells (HSPCs). Following transplantation into irradiated mice, the animals produced red blood cells (RBCs) with the EGFA domain displayed at the cell surface (EGFA-GPA RBCs). We assessed the potential of PCSK9 inhibitor-carrying RBCs as a long-term strategy to treat hyperlipidemia. We observed significant decreases in both plasma PCSK9 and LDL levels for at least 12 months. Interestingly, we also observed that the transplanted mice weighted significantly less than the control mice.

## Materials and Method

### Plasmids and generation of recombinant retroviruses

cDNA sequences of mouse EGFA and human GPA, National Center for Biotechnology Information (NCBI) reference sequences NC_000075.6 and NM_002099.6, respectively, were synthesized and then cloned into XhoI and EcoRI-cut XZ201 using a Thermo Scientific Rapid DNA ligation kit following the manufacturer’s protocol. The GFP-expressing, XZ201 plasmids were used for the production of murine stem cell virus (MSCV)-based retrovirus vectors to infect erythroid progenitors (Figure 1A). Therefore, positive transduction is indicated by GFP expression.

**Fig 1.**
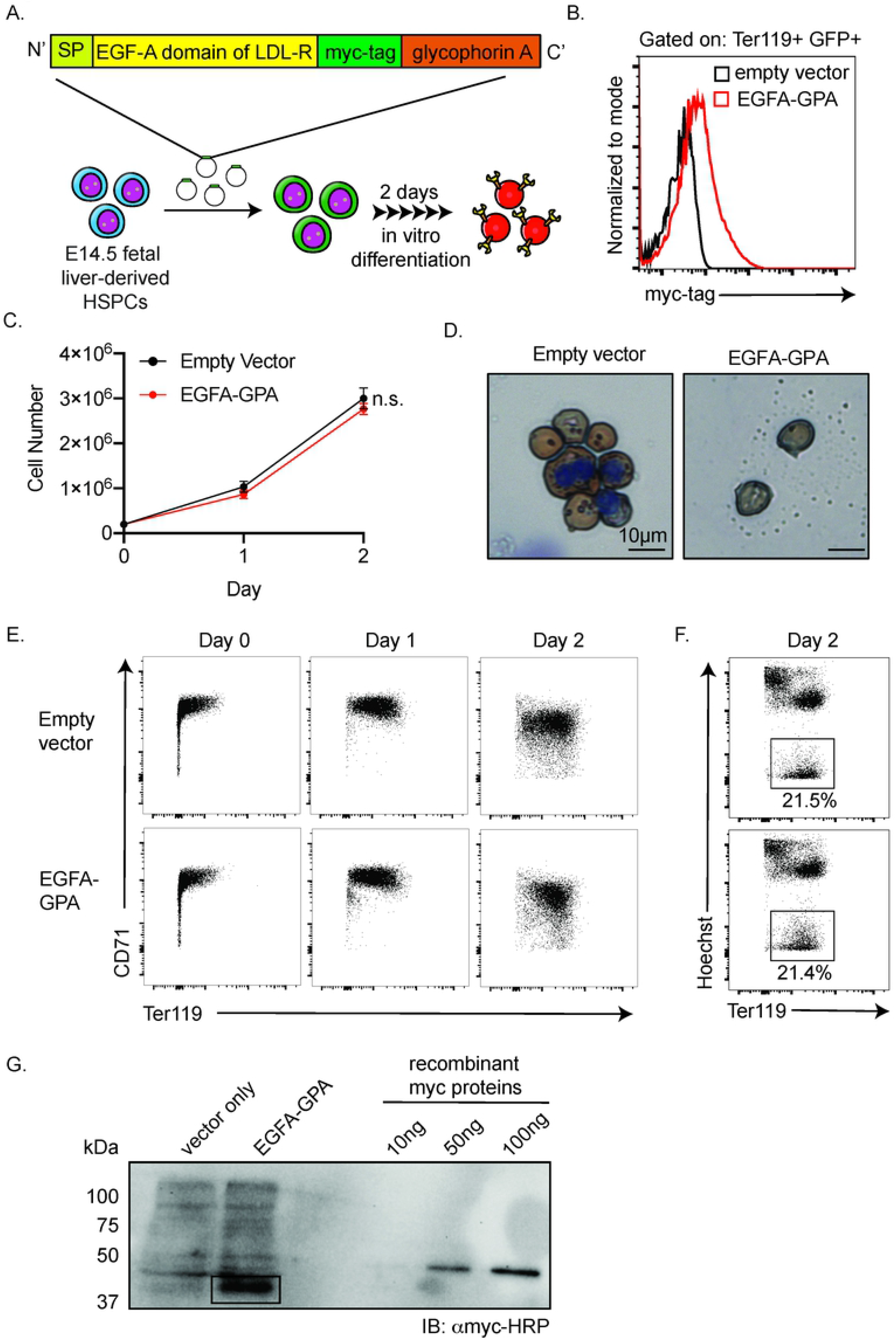
Reticulocytes produced in cell culture can be engineered to express on their surface the EGFA domain of LDLR without compromising their normal biological properties. (A) Design of the chimeric EGFA-GPA protein. Schematic depicting the generation of engineered reticulocytes produced by *in vitro* culture of murine fetal liver-derived HPSCs (SP = signal peptide) (B) Surface myc-tag expression of murine reticulocytes generated in culture that express empty vector (expressing GFP under the control of the CMV promoter to help indicate positive transduction in cells) or EGFA-GPA together with GFP, as evaluated by flow cytometry. (C) Proliferation of *in vitro*-differentiated erythroid cells (n = 3) (D) Benzidine-Giemsa staining of the resulting reticulocytes. (E) Representative flow cytometry evaluation of erythroid differentiation progression as indicated by surface expression of CD71 (transferrin receptor) and Ter119 (marker of mature erythroid cells) and (F) enucleation rate of erythroid cells as indicated by low Hoechst and high Ter119 staining. (G) Quantification of the number of EGF-A molecules per erythroid cell by western blot. Lysates from 1 million erythroid cells collected at day 2 terminal differentiation were analyzed. As quantification standard, we analyzed 10, 50, and 100 ng of full-length recombinant human c-Myc protein (molecular weight = 55 kD). Western blots were performed with anti-myc antibody. Calculated from the band’s density, each EGFA-GPA erythroid cell expresses ~9.36 million EGFA-GPA protein.

HEK293T cells were cultured in Dulbecco’s Modified Eagle Medium (DMEM) with 10% fetal bovine serum (FBS) in a humidified 5% CO2 atmosphere at 37°C. XZ201-based plasmids and pCLECO packaging plasmids were transformed into HEK293T cells with Fugene 6 (Promega) according to the manufacturer’s protocol. The medium was replaced after 8 hours, and retrovirus - containing supernatant was collected and filtered after 24 hours for transduction.

### Flow cytometric analyses and antibodies

All flow cytometric data were acquired on a Fortessa flow cytometer (BD Biosciences) and analyzed using Flowjo software (Tree Star). All stainings were carried out in FACS buffer (2mM EDTA and 1% FBS in phosphate-buffered saline/PBS) for 20 minutes on ice. Samples were washed twice with FACS buffer prior to flow analyses. The following are the antibodies used, all at 1:100 dilution: anti-myc tag-PE (Cell Signaling Technology, 3739), anti-mouse Ter119-APC (eBioscience, 17-5921-83), and anti-mouse CD71-PE (Affymetrix, 12-0711-83). Hoechst 33342 (Life Technologies, H1399) was used to visualize nuclei.

### Isolation, viral infection and culture of murine erythroid progenitors

Pregnant C57BL/6J mice at embryonic day 14.5 (E14.5) were euthanized by CO2 asphyxiation. Fetal livers were isolated from the embryos and suspended in cold PBS with 2% FBS and 100 μM EDTA. Mature RBCs in the cell suspension were lysed by incubation in sterile ammonium chloride solution (Stemcell) for 10 min at 4°C. Lineage-positive cells were magnetically depleted using the BD Pharmingen Biotin MouseLineage Panel (559971; BD Biosciences) and BD Streptavidin Particles Plus-DM (557812; BD Biosciences), per the manufacturer’s protocol. Lineage-negative fetal liver cells, enriched for erythroid progenitors, were then plated in 24-well plates at 100,000 cells per well, covered by 1 ml of virus containing supernatant, produced as described above, and 0.1% polybrene (Sigma Aldrich), and spin-infected at 500 g for 90 minutes at 25°C. Cells were then cultured in fresh erythroid maintenance medium (StemSpan-SFEM; StemCell Technologies) supplemented with 100 ng/ml recombinant mouse SCF (R&D Systems), 40 ng/ml recombinant mouse IGF1 (R&D Systems), 100 nM dexamethasone (Sigma), and 2 U/ml erythropoietin (Amgen) at 37°C for 24 hours.

To collect virally infected erythroid progenitors, after 24 hours in culture cells were sorted for GFP+ by flow cytometry and cultured in fresh erythroid differentiation medium (IMDM containing 15% (vol/vol) FBS (Stemcell), 1% detoxified bovine serum albumin (BSA; Stemcell), 500 μg/ml holo-transferrin (Sigma-Aldrich), 0.5 U/ml Erythropoietin (Epo; Amgen), 10 μg/ml recombinant human insulin (Sigma-Aldrich), and 2mM L-glutamine (Invitrogen) at 37°C for 48 hours. Cells in differentiation medium were counted manually with a hemocytometer at 0 hour, 24 hours, and 48 hours. After 48 h, erythroid cells were fixed on slides with cold methanol and stained with May-Grünwald– Giemsa (Sigma) and diaminobenzidine hydrochloride reagents (Sigma-Aldrich GS-500 and D-9015) for morphological analyses.

### Calculation of the number of chimeric proteins on murine RBCs

Western blotting on 1,000,000 in vitro - differentiated murine erythroid cells was conducted with anti-myc-tag (9B11) mouse monoclonal antibody (Cell Signaling no. 2040, 1:1,000 dilution). The copy number of recombinant human GPA protein per erythroid cell was derived using a linear signal intensity plot generated with 10, 50, and 100 ng of recombinant human c-Myc tagged proteins (55 kDa). As determined by the myc signal intensity in the western blot with an anti-myc antibody and as quantified by ImageJ, 1,000,000 murine EGFA-GPA cells contained ~855 ng EGFA-GPA protein. The molecular weight of EGFA-GPA is ~40,000 Da. Thus, each reticulocyte expresses ~9,000,000 copies of EGFA-GPA.

### Transplantation of transduced mouse fetal liver cells

A total of 1,000,000 virally infected murine hematopoietic stem/ progenitor cells cultured in erythroid maintenance medium, produced as described above, were collected and resuspended in 100μl PBS. These cells were then injected retro-orbitally into a C57BL/6J (Jackson Laboratory) mouse irradiated at 1050 rad, 1 day prior to transfer, in a Gammacell 40 irradiator chamber (Nordion International Inc.), and transplanted mice were monitored for 4 weeks before further analyses.

### Transfusion and in vivo survival of EGFA-GPA RBCs

A total of 200 μl blood from transplanted mice containing RBCs expressing the EGFA-GPA chimeric protein was collected into heparinized tubes (Fisher Scientific, 365965). The RBCs were washed twice in PBS and resuspended in PBS. Labeling was then carried out with 5 μM CellTrace Violet Dye for 20 minutes at room temperature (Life Technologies). 20% FBS in PBS was then added to RBCs for quenching the staining reaction. Violet-labeled RBCs were washed twice with PBS and resuspended in 200μl sterile PBS for intravenous injection into recipient mice. An equal volume of unmodified RBCs, similarly stained, served as a control. A drop of blood, ~20 μl, was collected into heparinized tubes by retro-orbital bleeding 1 h after transfusion at day 0 and every 3–4 days for 1 month as indicated in the text. These blood samples were washed once with FACS buffer and then subjected to staining with anti-Ter119-APC and anti-myc tag-PE for 20 minutes on ice. Samples were washed twice with FACS buffer prior to analyses on a FACS Fortessa flow cytometer for violet fluorescence and for GFP, Ter119, and myc signals.

### PCSK9 and LDL measurement

Blood samples (~100 μL) were collected by retro-orbital bleeding 3 h after transfusion at day 0 and at time points indicated in the text after bone marrow reconstitution. Plasma from these samples were isolated and stored at −80 °C after quick freezing with liquid nitrogen until all samples were collected for simultaneous analysis. The plasma samples were diluted (20 μL in 200 μL PBS) and then quantified for PCSK9 using a Mouse Proprotein Convertase 9/PCSK9 Quantikine ELISA Kit (R&D systems) and plasma LDL using Cholesterol Assay Kit - HDL and LDL/VLDL (Abcam #ab65390). The detailed procedures were provided with the kit.

## Results

### Engineered mouse RBCs that express the EGFA peptide at the cell surface

Our goal was to test the feasibility of a genetically engineered erythroid stem and progenitor cell therapy as a method to reduce LDL cholesterol. We generated retroviral constructs that can infect hematopoietic stem/progenitor cells (HSPCs) and allow expression of chimeric proteins on the surface of mature RBCs. We targeted glycophorin A (GPA) as the RBC membrane protein for the creation of a fusion protein, based on its abundance and RBC-specific expression. We genetically fused the EGFA-like domain of LDLR (EGFA) to the N-terminus of mature GPA to expose EGFA on surface of the RBC plasma membrane (Fig 1A). A myc-tag was inserted between the C-terminus of EGFA and N-terminus of GPA to facilitate detection. To mark all transfected cells and their progeny, the retroviral vector also contained a green fluorescent protein (GFP) under the control of the CMV promoter. We infected E14.5 mouse fetal liver derived HSPCs with these retroviruses as well as with an empty vector that expressed GFP as a control and cultured them *in vitro* to differentiate into reticulocytes. All terminally differentiated transfected and thus GFP+ reticulocytes expressed the chimeric GPA on their surface, as indicated by myc surface expression (Fig 1B). In this *in vitro* culture system, the EGFA-expressing cells underwent enucleation at a level similar to control cells. They showed similar CD71 and Ter119 surface expression, proliferation, and morphology compared to control cells, suggesting that the presence of the fusion does not disturb normal erythroid differentiation (Figure 1C-F). Each reticulocyte was estimated to express ~9 million copies of EGFA-GPA (Figure 1G; see Methods for calculation).

### Engineered mouse RBCs expressing EGFA peptides do not perturb erythropoiesis *in vivo*

Since our engineered erythroid progenitors underwent normal erythropoiesis *in vitro*, we next produced engineered RBCs *in vivo* by transplanting sub-lethally irradiated mice with fetal liver hematopoietic stem/ progenitor cells (HSPCs) infected by the retrovirus expressing both GFP and EGFA-GPA. Complete blood counts (CBCs) and myc surface expression (indicating the expression of the EGFA-GPA chimera) from one batch of EGFA-GPA-transplanted mice were followed over time (Fig 2B). Compared to control 7-week-old female mice, the blood parameters of the transplanted mice were within the normal range reported in the Mouse Phenome Database on the Jackson Laboratory website. Moreover, all of the GFP+ cells, the progeny of the transfected HSPCs, expressed myc on their surface (Fig 2C). In a separate batch of mice transplanted with HSPCs expressing EGFA-GPA, 7.59 ± 1.34% of the RBCs were myc+ 4-weeks after bone marrow reconstitution (±S.E.M.; n = 12). There was no noticeable change in morphology and apoptotic marker in the engineered RBCs compared to controls (Figs 2D, E). To determine whether the introduction of EGFA-GPA impacts the circulatory half-life of RBCs, we stained isolated RBCs from the transplanted mice with the CellTrace Violet Dye prior to transfusion into normal recipients to monitor the total population of transfused RBCs. Cells expressing EGFA-GPA were monitored by the green fluorescent protein (GFP) signal (from the GFP expression cassette in the retroviral vector; Fig 2F). The circulatory lifespan of the transfused control and EGFA-GPA RBCs is similar to that of normal mouse RBCs (Fig 2F).

**Fig 2.**
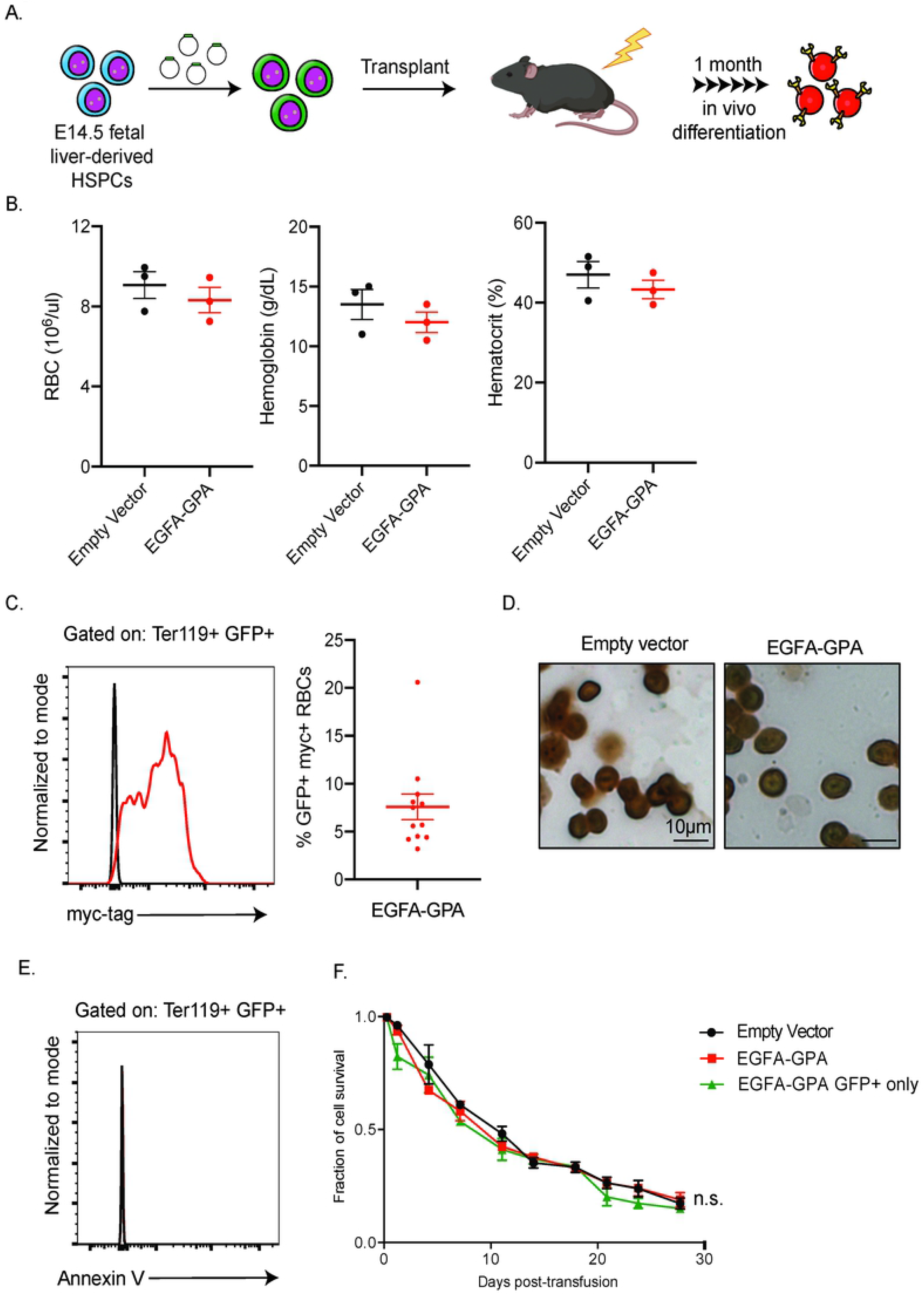
*In vivo* production of mature RBCs expressing EGFA-GPA. (A) Schematic depicting the generation of engineered mature RBC produced by transplanting genetically modified HSPCs. (B) Complete blood counts of 8-week-old female C57BL/6J mice subjected to bone marrow transplantation with progenitor cells expressing vector or EGFA-GPA and bled at 1-month post-transplantation. (n = 3/group, mean ± SEM) (C) At 1-month post-transplantation blood was analyzed by flow cytometry for the percentage of cells that were myc positive (right). As plasmids used to engineer the transplanted HSPCs contain GFP under the control of the CMV promoter to help indicate positive transduction, thus Ter119+ erythroid cells derived from the transplanted HSPCs were also GFP+ and were then analyzed for myc expression. (n = 12, mean ± SEM). (D) Benzidine-Giemsa and (E) Annexin V staining of the resulting RBCs. (F) Circulatory half-life of the transfused RBCs. 100 μl of blood from transplanted mice with ~8% of their RBCs expressing vector only or EGFA-GPA were stained with violet-trace dye and transfused into recipient mice. The fraction of transfused RBCs in recipients was analyzed by flow cytometry at the indicated time points. The violet-trace dye represents the total population of transfused RBCs, of which only ~8% are GFP+ and express the exogenous chimeric protein, while the GFP signal represents only the 8% of the transfused RBCs expressing the EGFA-GPA. (n = 3/group, mean ± SEM).

### Mice transplanted with HSPCs expressing EGFA-GPA maintained lower plasma PCSK9 and LDL levels for at least one year

Mice expressing engineered RBCs were fed a normal diet for 12 months, and their weight, plasma LDL and PCSK9 were measured every 3 months. Over the course of this period, the numbers of EGFA-GPA RBCs, as detected by GFP+ myc+ RBCs, in transplanted mice, remained around 3-6% of total RBCs (Fig 3A; Supplemental Fig 1). We observed significant decreases in plasma LDL and PCSK9 levels in EGFA-GPA transplanted mice. Engineered RBCs thus sequester circulatory PCSK9 (Figs 3B and C). Lower levels of plasma LDL in these mice are correlated with a significant reduction in body weight gain over the ensuing 12 months (Fig 3D).

**Fig 3.**
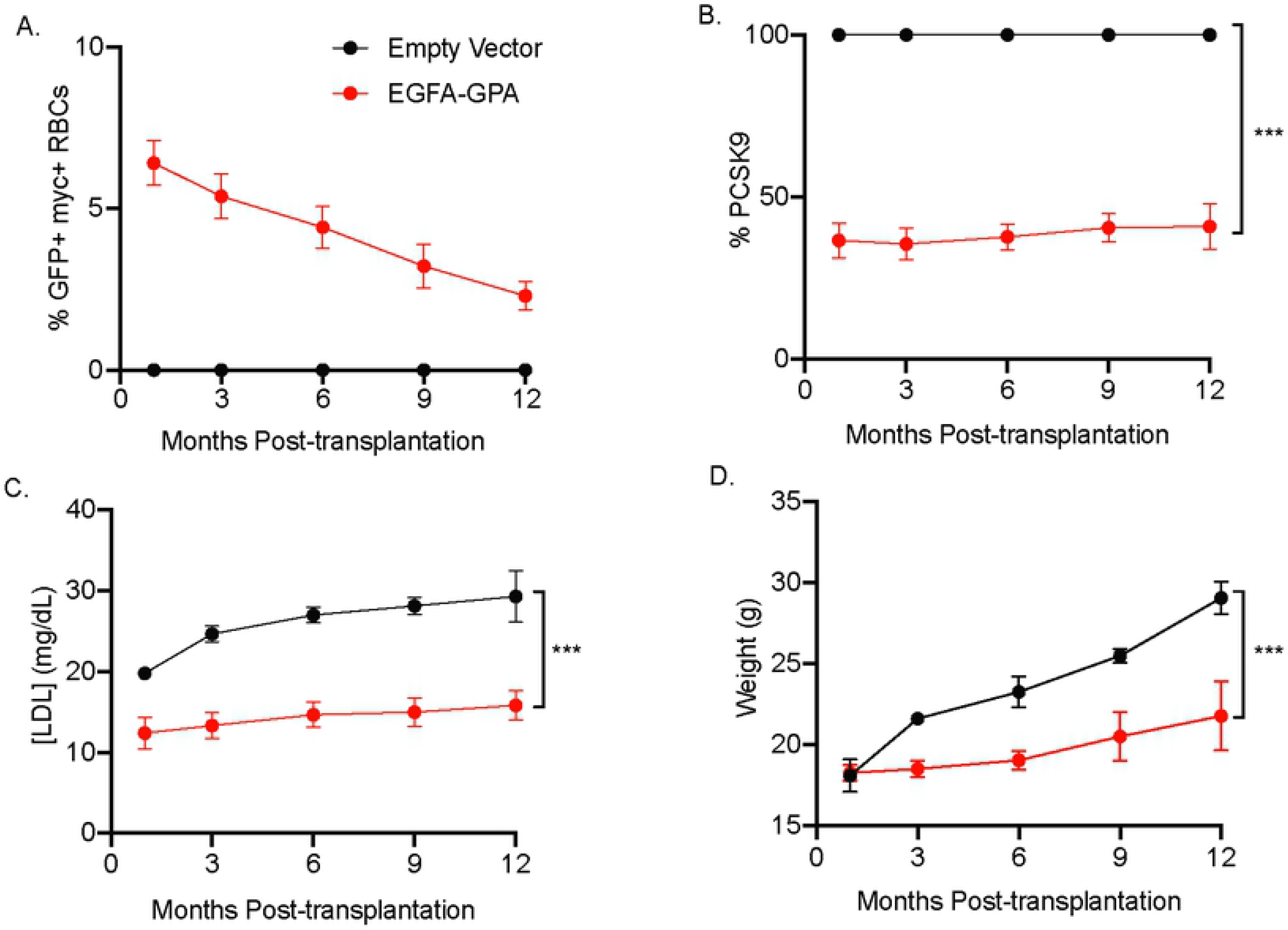
EGFA-GPA – expressing mice maintain lower plasma LDL level and weight for the remainder of their lifetime. (A) Long-term expression of myc+ cells (n = 12, mean ± SEM) in mice transplanted with HSPCs expressing the Glycophorin A-EGF-A chimeric protein. (B) Plasma PCSK9 and (C) LDL levels of transplanted mice were quantified via ELISA assay at the indicated time points. Engineered RBCs act as a sink in capturing circulating PCSK9 molecules, inhibiting LDLR degradation that resulted in lower plasma LDL concentration. (n = 8/group, mean ± SEM). (D) Mice expressing EGFA-GPA RBCs weigh significantly less than control mice throughout their observed lifespan (n = 8/group, mean ± SEM).

### EGFA-GPA transplanted mice resist high-fat diet-induced obesity

To better mimic the type of diet that causes obesity, we fed mice a high-fat diet (~60% fat chow) for 8 weeks, and measured their weight, plasma LDL, and PCSK9 biweekly. The high-fat diet for mice also caused elevated LDL levels that are more similar to humans. The numbers of EGFA-GPA RBCs in transplanted mice, as indicated by GFP+ myc+ RBCs, remained steady at about 9-12% (Fig 4A; Supplemental Fig 2). Circulating PCSK9 levels in mice that carry EGFA-GPA RBCs were significantly lower than in controls across all time points, although PCSK9 levels did gradually increase over these 8 weeks (Fig 4B). Unexpectedly we did not observe a significant decrease in plasma levels of LDL (Fig 4C). Nonetheless, EGFA-GPA transplanted mice were still able to resist high-fat diet induced weight gain and maintained their weight for 8 weeks. This established the efficacy of the engineered RBCs in preventing diet-induced obesity (Fig 4D). Hematoxylin and eosin staining of various organs including heart, kidney, liver and spleen showed no abnormalities. We conclude that the presence of EGFA-GPA RBCs does not have obvious side effects despite their protection against obesity (Fig 4E).

**Fig 4.**
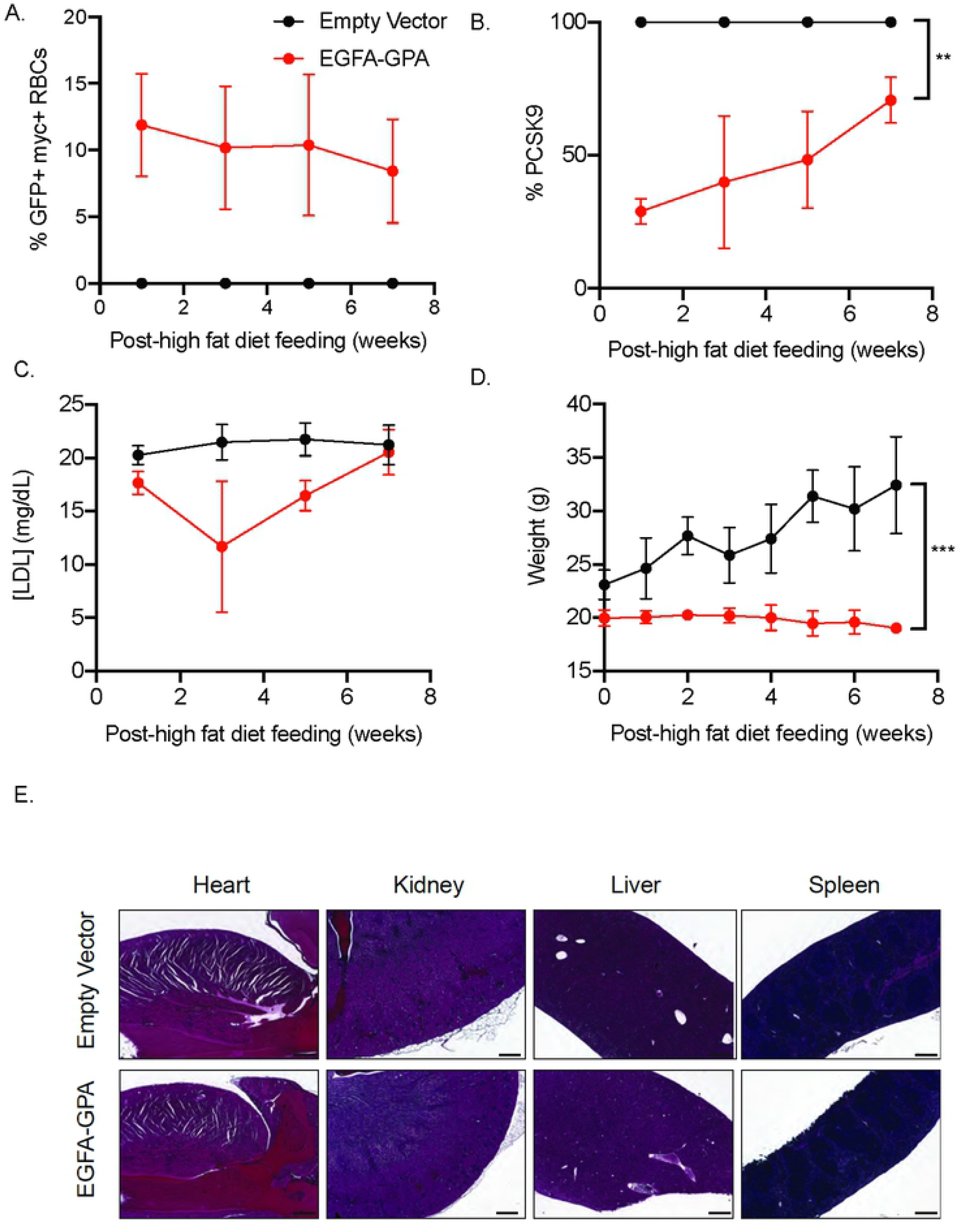
Hematopoietic transplantation of engineered RBCs prevent weight gained in mice fed with a high-fat diet but unable to lower LDL levels. Mice were placed on a high-fat diet 4 weeks after transplantation with HSPCs expressing the EGFA-GPA chimeric protein and monitored biweekly. (A) Long-term expression of myc+ cells (n = 3, mean ± SEM). (B) Plasma PCSK9 and (C) LDL levels of transplanted mice were quantified via ELISA assay at indicated time points. (n = 3/group, mean ± SEM). (D) Mice expressing EGFA-GPA RBCs weigh significantly less than control mice (n = 3/group, mean ± SEM). (E) Hematoxylin and eosin staining of various organs: heart, kidney, liver and spleen.

## Discussion

We showed that mouse RBCs, engineered to expressing on their surface a peptide PCSK9 inhibitor, can deplete circulatory PCSK9 by acting as a systemic sink. This depletion results in a persistent reduction of plasma LDL levels. Animals that carry EGFA-GPA RBCs upon receiving modified hematopoietic stem cells, were resistant to diet-induced obesity. Transplantation of autologous, genetically modified HSPC that express an EGFA-GPA chimeric protein, could therefore be a practical and one-shot preventative treatment for cardiovascular disease and obesity in humans, especially those with a genetic predisposition to high cholesterol levels.

Selective bone-marrow depletion techniques that target and kill specific hematopoietic progenitors, especially HSPCs, avoid complete myeloablation. This means that patients can undergo bone-marrow transplantation without chemotherapy or radiotherapy and avoid the toxic side effects that come with such treatments [19, 20]. We envision that this approach can provide much lasting effect on lowering LDL and preventing obesity. We observed that the percentage of engineered RBC expressing the EGFA-GPA decreases overtime in transplanted cohort. However, the levels of PCSK9 and LDL did not correlate with the decreasing EGFA-GPA RBC levels. This result suggested that we potentially could reach similar decrease of PCSK9 and LDL levels at lower percentage of RBC reconstitution (<2%). This could also explain the low variation in PCSK9 levels of transplanted cohort despite the variation in the engineered RBC expression. We did not test this hypothesis due to the technical difficulties in titrating the percentage of engineered RBC expressed in transplanted mice. We also envision that more potent inhibitors than the EGFA peptide might achieve even greater reductions of plasma LDL levels and with fewer engineered RBCs. For example, one can covalently attach EGFA peptide variants with enhanced affinity for PCSK9 act as a systemic sink, or anti-PCSK9 camelid-derived single domain antibody fragments onto the glycophorin A protein [21].

It is an intriguing finding that we observed a reduction in weight gain and resistance to diet induced obesity in the EGFA-GPA mice. Since PCSK9 also enhances degradation of vLDLR expressed in adipocytes, we actually expected the opposite result. Indeed, PCSK9 knock-out mice show increased uptake of chylomicrons and vLDL accumulation in visceral fat, adipocytes, heart and muscles, i.e. in tissues where vLDLR is expressed predominantly. However, PCSK9 knock-out mice are not prone to obesity [22]. We suggest that using the EGFA-GPA modified red blood cells, which persistently deplete only the circulatory PCSK9, may allow PCSK9 to continue to function normally in the liver in autocrine and paracrine manner. This may have affected lipid metabolism differently in comparison with complete ablation of PCSK9 in knock-out mice. Similarly, antibody treatment allows for inhibition of both circulating and liver PCSK9. Hence, we speculate that the decrease of weight gain is attributed to the inhibition of only circulating PCKS9 that is uniquely achieved in transplanted mice.

The depletion of only the circulating PCSK9 could also explain the reason why this approach unable to lower LDL levels when mice fed with high-fat diet. Mice lipoprotein profile is different than human, in which majority of plasma cholesterol are in the form of HDL while in human LDL accounts for majority of total plasma cholesterol. The high-fat diet caused significant increase of LDL in mice to the levels that are more similar to human. Our engineered RBC approach robustly decreased the PCSK9 levels in plasma, however, it may fail to significantly decrease the levels of PCSK9 in the extracellular spaces on the liver. Since liver accounts for the majority of LDL clearance, inhibition of only the circulating PCSK9 activity may not sufficient to cause significant increase of LDL uptake, hence we observe very little to no decrease of LDL levels in transplanted mice. We did still observe weight difference in transplanted vs. control cohort even to a greater degree, suggesting an unexplored function of circulating PCSK9 in lipid accumulation in peripheral tissues.

We sought to develop a more practical treatment to lower LDL via a single transfusion of engineered hematopoietic stem cells to produce PCSK9 inhibitor-carrying RBCs. The depletion of circulating PCSK9 lowered LDL levels in transplanted mice, however, failed to effectively treat hypercholesterolemia under high-fat diet. Unexpectedly, we observed significant difference in total body weight of transplanted vs. control cohort. Since PCSK9 has a variety of functions outside the liver, not all of which are fully understood, the EGFA-GPA mice reported here could serve as a model to systematically explore the function of circulatory PCSK9 in the pathogenesis of diabetes, inflammation, sepsis, viral infections, and especially, obesity. With obesity and cardiovascular disease as main causes of death worldwide, a safe, effective and permanent therapy to inhibit PCSK9 can improve health span and quality of life. This preliminary study suggested a potential of using red blood cells genetically modified as circulating PCSK9 inhibitors to treat hyperlipidemia and obesity.

**Supplementary Fig 1.** Individual values corresponding to data depicted in Figure 3.

**Supplementary Fig 2.** Individual values corresponding to data depicted in Figure 4.

## Acknowledgments

We thank members of the H.F.L. laboratory, especially Tony Chavarria and Ferenc Reinhardt for mouse husbandry. This work was supported by Defense Advanced Research Projects Agency Contract HR0011-12-2-0015 (to H.L.P. and H.F.L.) and grants from the Howard Hughes Medical Institute International Student Research Fellowship and the Siebel Scholarship (to N.P.).

## Disclosures

H.F.L. and H.L.P. served as scientific advisors and have/had equity in Rubius, a biotechnology company that seeks to exploit engineered red blood cells as therapeutics but does not provide financial support for the technology described in this paper.

## References

1. Naci H, et al. Comparative Benefits of Statins in the Primary and Secondary Prevention of Major Coronary Events and All-Cause Mortality: A Network Meta-Analysis of Placebo-Controlled and Active-Comparator Trials. European Journal of Preventive Cardiology. 2013; doi:10.1177/2047487313480435.

2. Abifadel M, et al. Mutations in PCSK9 Cause Autosomal Dominant Hypercholesterolemia. Nature Genetics. 2003; doi:10.1038/ng1161.

3. Cohen J, et al. Low LDL Cholesterol in Individuals of African Descent Resulting from Frequent Nonsense Mutations in PCSK9. Nature Genetics. 2005; doi:10.1038/ng1509.

4. Seidah NG and Annik P. The Biology and Therapeutic Targeting of the Proprotein Convertases. Nature Reviews Drug Discovery. 2012; doi:10.1038/nrd3699.

5. Zhang Y, et al. Identification of a Small Peptide That Inhibits PCSK9 Protein Binding to the Low Density Lipoprotein Receptor. Journal of Biological Chemistry. 2014; doi:10.1074/jbc.M113.514067.

6. Seidah NG, et al. PCSK9: A Key Modulator of Cardiovascular Health. Circulation Research. 2014; doi:10.1161/CIRCRESAHA.114.301621.

7. Demers A, et al. PCSK9 Induces CD36 Degradation and Affects Long-Chain Fatty Acid Uptake and Triglyceride Metabolism in Adipocytes and in Mouse Liver. Arteriosclerosis, Thrombosis, and Vascular Biology. 2015; doi:10.1161/ATVBAHA.115.306032.

8. Levy E, et al. PCSK9 Plays a Significant Role in Cholesterol Homeostasis and Lipid Transport in Intestinal Epithelial Cells. Atherosclerosis. 2013; doi:10.1016/j.atherosclerosis.2013.01.023.

9. Bergeron N, et al. Proprotein Convertase Subtilisin/Kexin Type 9 Inhibition. Circulation. 2015; doi:10.1161/CIRCULATIONAHA.115.016080.

10. Iskowitz M. Can the Third PCSK9 Drug Succeed Where the First Two Failed? MM&M - Medical Marketing and Media, 6 Feb. 2020. www.mmm-online.com, https://www.mmm-online.com/home/channel/features/can-the-third-pcsk9-drug-succeed-where-the-first-two-failed/.

11. Qiurong D, et al. Permanent Alteration of PCSK9 With In Vivo CRISPR-Cas9 Genome Editing. Circulation Research. 2014; doi:10.1161/CIRCRESAHA.115.304351.

12. Shi J, et al. Engineered Red Blood Cells as Carriers for Systemic Delivery of a Wide Array of Functional Probes. Proceedings of the National Academy of Sciences. 2014; doi:10.1073/pnas.1409861111.

13. Pishesha N, et al. Engineered Erythrocytes Covalently Linked to Antigenic Peptides Can Protect against Autoimmune Disease. Proceedings of the National Academy of Sciences. 2017; doi:10.1073/pnas.1701746114.

14. Huang N-J, et al. Genetically Engineered Red Cells Expressing Single Domain Camelid Antibodies Confer Long-Term Protection against Botulinum Neurotoxin. Nat Commun. 2017; doi:10.1038/s41467-017-00448-0.

15. Mamcarz E, et al. Lentiviral Gene Therapy Combined with Low-Dose Busulfan in Infants with SCID-X1. N Engl j Med. 2019;380(16):1525–1534. doi:10.1056/NEJMoa1815408.

16. Ferrua F, et al. Lentiviral haemopoietic stem/progenitor cell gene therapy for treatment of Wiskott-Aldrich syndrome: interim results of a non-randomised, open-label, phase 1/2 clinical study. Lancet Haematol. 2019;6(5):e239–e253. doi:10.1016/S2352-3026(19)30021-3.

17. Zeng J, et al. Therapeutic base editing of human hematopoietic stem cells. Nat Med. 2020; 26(4):535–541. doi:10.1038/s41591-020-0790-y.

18. Gennery AR, Albert MH, Slatter MA, Lakester A. Hematopoietic Stem Cell Transplantation for Primary Immunodeficiencies. Front Pediatr. 2019; 7:445. doi:10.3389/fped.2019.00445.

19. Chhabra A, et al. Hematopoietic Stem Cell Transplantation in Immunocompetent Hosts without Radiation or Chemotherapy. Science Translational Medicine. 2016; doi:10.1126/scitranslmed.aae0501.

20. Czechowicz A, et al. Selective Hematopoietic Stem Cell Ablation Using CD117-Antibody-Drug-Conjugates Enables Safe and Effective Transplantation with Immunity Preservation. Nature Communications. 2019; doi:10.1038/s41467-018-08201-x.

21. Weider E, et al. Proprotein Convertase Subtilisin/Kexin Type 9 (PCSK9) Single Domain Antibodies Are Potent Inhibitors of Low Density Lipoprotein Receptor Degradation. Journal of Biological Chemistry. 2016; doi:10.1074/jbc.M116.717736.

22. Roubtsova A, et al. Circulating Proprotein Convertase Subtilisin/Kexin 9 (PCSK9) Regulates VLDLR Protein and Triglyceride Accumulation in Visceral Adipose Tissue. Arteriosclerosis, Thrombosis, and Vascular Biology. 2011; doi:10.1161/ATVBAHA.110.220988.

